# Distractor inhibition contributes to retroactive attentional orienting within working memory: Evidence by lateralized alpha oscillations in the EEG

**DOI:** 10.1101/235069

**Authors:** Daniel Schneider, Anna Barth, Henrike Haase, Clayton Hickey, Edmund Wascher

**Affiliations:** Leibniz Research Centre for Working Environment and Human Factors, TU Dortmund; Center for Mind/Brain Sciences (CiMeC), University of Trento

**Keywords:** Working memory, attention, retro-cue, alpha oscillations, selective forgetting

## Abstract

Shifts of attention within mental representations based on retroactive cues (retro-cues) facilitate performance in working memory tasks. It was suggested that this retro-cue benefit is related to the concentration of working memory resources on a subset of representations, thereby improving storage and retrieval at the cost of non-cued items. However, the attentional mechanisms underlying this updating of working memory representations remain unknown. Here, we present EEG data for distinguishing between target enhancement and distractor suppression processes in the context of retroactive attentional orienting. Therefore, we used a working memory paradigm with retro-cues indicating a shift of attention to either a lateralized or non-lateralized item. There was an increase of posterior alpha power contralateral compared to ipsilateral to the irrelevant item when a non-lateralized mental representation was cued and a contralateral suppression of posterior alpha power when a lateralized item had to be selected. This suggests that both inhibition of the non-cued information and enhancement of the target representation are important features of attentional orienting within working memory. By further presenting cues to either remember or to forget a working memory representation, we give a first impression of these retroactive attentional sub-processes as two separable cognitive mechanisms.

## 1. Introduction

Goal-directed behavior under changing environmental conditions requires to memorize only those mental representations over the short-term that are relevant for current action. This entails shifting the focus of attention within short-term or working memory representations and focusing on certain representations at the expense of no-longer relevant information. By means of event-related parameters in the EEG, the current study investigated to what extent excitatory and inhibitory processes contribute to this orienting of attention on the level of working memory.

So-called retroactive cuing (or retro-cue) paradigms are suited for studying the allocation of attention within working memory. It could be shown that comparable to cues causing a shift of attention toward a subset of to-be-presented stimuli in a memory array, also those presented after encoding facilitate performance in visuo-spatial working memory paradigms (Griffin & Nobre, 2003; Oberauer, 2002). Here, we focus on the sub-processes that underlie attentional selection within visuo-spatial working memory representations. When speaking of ‘selection’, there are two potentially relevant mechanisms: Attentional templates for target features lead to a top-down bias on the firing of neurons coding for these features, thereby leading to a representational advantage in the course of visual processing (i.e. ‘target enhancement’). Additionally, an attentional template might also function by turning down the response of neurons coding for non-relevant features. This can be mediated by top-down inhibitory mechanisms or by automatic lateral inhibition from the attended feature (or both; Desimone, 1998; Desimone & Duncan, 1995).

For attentional selection in the course of perception, it could already be shown that both target enhancement and distractor inhibition play a prominent role. Hickey, DiLollo and McDonald (2009) studied attentional sub-processes in visual search based on lateralized responses in the EEG. As our visual system is retinotopically organized to large extents, the authors presented either distractor or target items on lateral positions in a visual search display. A prominent component in the event-related potential (ERP) of the EEG, the N2 posterior contralateral or N2pc (Luck & Hillyard, 1994a, 1994b), was thereby divided into a posterior contralateral negativity indexing target enhancement and a posterior contralateral positivity reflecting distractor inhibition.

Retroactive attentional orienting within working memory was also shown to elicit retinotopically organized asymmetries in the EEG, with studies usually focusing on a decrease in posterior oscillatory power in the alpha frequency range contralateral to selected visuo-spatial representations (Myers, Walther, Wallis, Stokes, & Nobre, 2015; Poch, Capilla, Hinojosa, & Campo, 2017; Poch, Carretie, & Campo, 2017; Schneider, Mertes, & Wascher, 2015, 2016). We thus made use of the spatial separation of target and distractor stimuli and combined it with two retro-cue based working memory tasks in order to separate target enhancement and distractor inhibition processes reflected in posterior alpha asymmetry. In Experiment 1, we presented a retro-cue indicating the relevant memory array item and further used a probe presented at the time of the retro-cue as a behavioral control condition. If retroactive attentional selection is based on target enhancement, we should observe a contralateral decrease in posterior alpha power with a lateralized target as a sign for target enhancement, while an increase in alpha power contralateral to the non-cued item would point toward distractor inhibition on the level of working memory. In Experiment 2, we compared these attentional sub-processes between retro-cue conditions with cues indicating either the target (i.e., remember cues) or distractor positions (i.e., forget cues). This was done in order to ascertain if a potential contribution of inhibitory mechanisms to retroactive attentional orienting is related to an active cognitive process (i.e., selective forgetting) or to lateral inhibition by means of the attentional template for the target item. We hypothesized that if selective forgetting contributes to the retro-cue benefit in working memory, then forget cues should lead to a facilitation of distractor inhibition processes relative to remember cues. However, if forget cues are reframed to function as remember cues, there should be a general facilitation of attentional orienting in favor of the remember cues.

## 2. Materials and Methods

### 2.1. Participants

Twenty participants (9 females; M(age)=25.15 years, SD=3.58 years, range: 19-30 years) took part in Experiment 1 and 16 participants took part in Experiment 2 (7 females; M(age)=25.06 years, SD=3.33 years, range: 19-29 years). None of the participants in Experiment 1 participated in Experiment 2. All participants were right-handers. As shown by means of a screening questionnaire, none of them reported any known neurological or psychiatric diseases and had normal (non-corrected) vision. Participation was rewarded by a payment of 10 € per hour or course credit. All participants gave their informed consent for participation after receiving written information about the study’s purpose and procedure. The studies were run in accordance with the Declaration of Helsinki and approved by the local ethics committee at the Leibniz Research Centre for Working Environment and Human Factors.

### 2.2. Stimuli and procedure

The experiments were run on a 22-inch CRT monitor (100 Hz) with a viewing distance of 150 cm and a display resolution of 1024 × 768 pixels. Stimulus presentation was done by means of a ViSaGe MKII Stimulus Generator (Cambridge Research Systems, Rochester, UK). Throughout the whole trial, a black fixation cross was presented on a dark gray background (luminance of 15 cd/m^2^). Participants were instructed to always fixate. This was furthermore controlled by an SMI Red 500 eye-tracking device (SensoMotoric Instruments, Teltow, Germany) measuring eye-movements at a frequency of 120 Hz. Prior to the beginning of the experiment and after each experimental block (see below), eye tracking was calibrated by measuring eye movements while presenting a gray circle (25 cd/m^2^) on a dark gray background (10 cd/m^2^) that successively moved to five different positions. We defined a 2° × 2° (visual angle) area around the central position of the screen and fixation control during the experiment was then carried out by only starting a trial (i.e. presenting the initial memory array), when participants fixated for at least 40 ms within a 100 ms interval prior to the memory array. The intertrial interval was set to 500-1000 ms (randomized), but was extended by on average 409 ms (SD=198.72 ms) due to the fixation control procedure.

#### 2.2.1. Experiment 1

The memory array was composed of three bars (0.1 by 1° of visual angle; see figure 1) presented with a random orientation, under the premise that orientation had to differ between the bars by at least 15°. The bars were aligned on a hypothetical circle with 1.5° radius and presented at 60° (right), 180° (bottom) and 300° (left). Thus, one bar was presented below fixation (i.e. on the horizontal median). The other two bars were presented above fixation and placed to the left and right side. One of these lateralized bars was presented in gray with 25 cd/m^2^. The two remaining bars (lateralized position or central position below fixation) were also presented with 25 cd/m^2^, but either in color red (CIE: 0.566, 0.376, 0.25) or blue (CIE: 0.168, 0.131, 0.25). Accordingly, only one of the two lateralized items was colored, while the contralateral bar was always presented in gray (see figure 1). While the gray bar was never relevant for task performance, the red and blue bars were potentially relevant until presentation of the subsequent retro-cue or memory probe. The memory array was presented for 200 ms and followed by the retro-cue or a memory probe with an inter-stimulus interval of 800 ms. In the retro-cue condition, the cue was presented as an enlarged fixation cross and indicated the relevant memory item by color (red or blue). In 2/3 of all retro-cue trials, the cued item was presented left or right of fixation (i.e. target lateralized condition). In the remaining trials, the retro-cue pointed toward the central item below the fixation cross, indicating one lateralized working memory representation as irrelevant (i.e. distractor lateralized condition). This condition was presented in only 1/3 of the retro-cue trials to present the target equally frequent at each memory array position. The retro-cue was presented for 100 ms and followed by a further delay interval of 900 ms. Afterwards, a randomly oriented black bar (0 cd/m^2^) was presented at fixation and had to be matched in orientation to the previously cued item by moving the computer mouse. This memory probe remained presented for 3000 ms. Participants were instructed to press the left computer mouse button when they considered their orientation adjustment as appropriate.

**Figure 1.**
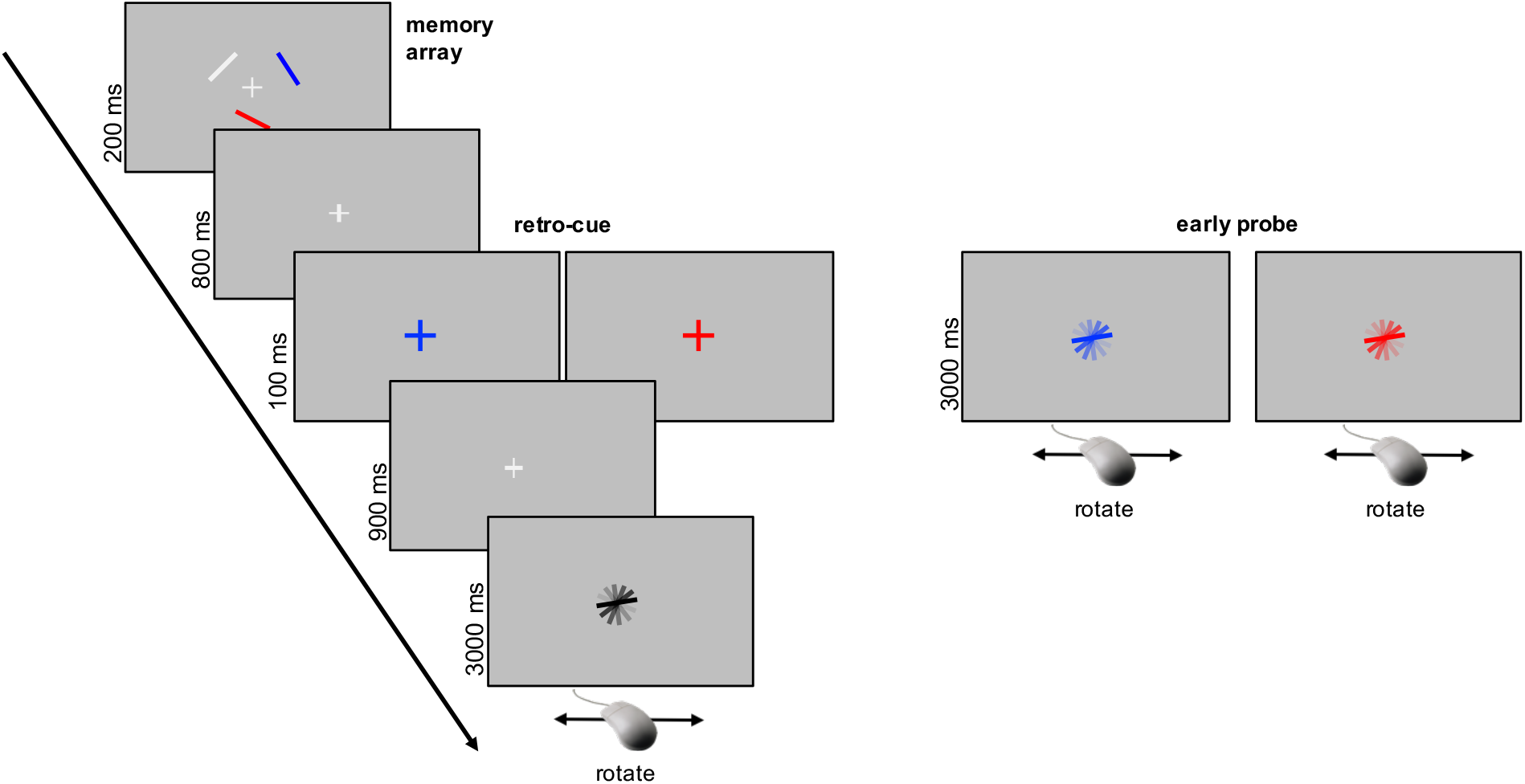
Experimental design. For both Experiment 1 and 2, each trial featured an initial memory array containing two potentially relevant randomly oriented bars (blue and red) and one irrelevant bar (gray). The irrelevant bar was always presented on a lateral position. In Experiment 1, a subsequent retro-cue or early probe indicated either the lateralized or non-lateralized (bottom) stimulus as relevant for the orientation adjustment task. In Experiment 2, the retro-cue indicated either the item to remember or to forget. No early probe condition was implemented in Experiment 2.

We furthermore included an ‘early probe’ condition in 50% of the trials (see figure 1) as a control condition for estimating the retro-cue benefit in the working memory task. In this condition, the retro-cue was replaced by the onset of the memory probe display that again remained visible for 3000 ms. Because early probes and retro-cues were presented at the same point in time, there was no difference in the time of working memory retrieval between these conditions. This has the advantage that retro-cue benefits referred to the early probe condition cannot be related to time-based decay of the working memory representations (Souza & Oberauer, 2016). The cued item was indicated by presenting the rotatable memory probe in either red or blue. The proportion of the target vs. distractor lateralized conditions was the same as in the retro-cue condition. Overall, the experiment consisted of 960 trials divided into 8 blocks of trials. A short break of two minutes was made between the blocks in order to prevent fatigue in the course of the experiment.

#### 2.2.2. Experiment 2

This experiment was run in order to investigate if inhibitory attentional sub-processes on the level of working memory are related to lateral inhibition related to the target information or to volitional or active forgetting of the irrelevant contents. While we did not include an ‘early probe’ condition, as this would have entailed a very long experiment, the other experimental parameters were similar to Experiment 1. The rate of memory items cued as targets was equally frequent for the three positions. However, the meaning of the retro-cue was varied in a block-wise fashion. It could indicate the color of either the to-be-remembered working memory item (‘remember cue’ condition) or the to-be-forgotten item (‘forget cue’ condition).

The order of the experimental blocks with remember vs. forget cues was counterbalanced across participants. Retro-cue meaning was changed after 4 experimental blocks (i.e. when half of the experiment was over). Thus, the experiment consisted of 8 blocks with 960 trials overall. A short break of two minutes was made between the blocks in order to prevent fatigue in the course of the experiment.

### 2.3. EEG recording and preprocessing

EEG was recorded with a frequency of 1000 Hz by means of a NeurOne Tesla AC-amplifier (Bittium Biosignals Ltd, Kuopio, Finland), based on 64 Ag/AgCl passive electrodes (Easycap GmbH, Herrsching, Germany) affixed across the scalp according to the extended 10/20 System. A 250 Hz low-pass filter was used during recording. The ground electrode was set to midline electrode AFz. The reference electrode during recording was at FCz. Impedance was kept below 10kΩ during recording.

We used MATLAB® and EEGLAB (Delorme & Makeig, 2004) for analyzing the EEG data. A 0.5 Hz high-pass and 30 Hz low-pass filter were applied prior to re-referencing the data to the average signal of all channels (average reference) and rejecting bad channels. Channel rejection was based on an absolute threshold limit of 5 SD regarding the kurtosis measure for each channel. Data epochs were then created beginning 1000 ms before and ending 3000 ms after memory array presentation. Independent component analysis (ICA) was run on every second epoch in the dataset and ADJUST (Mognon, Jovicich, Bruzzone, & Buiatti, 2010) was used to detect ICs related to eye blinks, vertical and horizontal eye movements and generic data discontinuities. The IC weights of the dataset reduced to every second trial were then transferred to the whole dataset and the selected ICs were removed from the signal. Additionally, we computed single dipoles for each IC by means of a boundary element head model (Fuchs, Kastner, Wagner, Hawes, & Ebersole, 2002), and also excluded ICs with a dipole solution with more than 40% residual variance. This procedure was followed by an automatic trial rejection procedure implemented in EEGLAB (threshold limit: 1000 µV, probability threshold: 5 SD, Max. % of trials rejected per iteration: 5%). These preprocessing steps led to the rejection of 165 trials on average (SD=51.97) in Experiment 1 and 177 trials on average in Experiment 2 (SD=51.46).

## 3. Results

### 3.1. Behavioral data

#### 3.1.1. Experiment 1

The raw angular errors were calculated by subtracting the orientation set for the memory probe from the original orientation of the cued item in the memory array. Therefore, the largest possible angular error was 90°. As a further parameter for assessing working memory accuracy, we calculated the standard deviation (SD) of the raw error within each experimental condition. These values were adjusted for circular data by means of the CircStat Toolbox for MATLAB® (Berens, 2009). Regarding parameters reflecting the speed of response, we chose to measure both the onset of the computer mouse movement and the time of the button press. While the time point of the button press should also differ as a function of the difference in orientation between the cued item and the memory probe, the onset of computer mouse movement should provide a reliable measure of the time required for response preparation. Only trials with a mouse button press were included in the behavioral analyses. The rate of button presses across trials was 99.81% on average (SD=0.29%). Cohen’s d_z_ for dependent measures is used as an indicator for effect size.

No difference in behavioral performance between the retro-cue and the early probe conditions appeared regarding the raw error, *t*(19)=−0.233, *p*=0.819, d_z_=−0.052, and SD of the raw error, *t*(19)=−0.911, *p*=0.374, d_z_=−0.204, indicating that in our sample the retro-cue did not lead to an increase in the overall accuracy of the recalled working memory representations (see figure 2A). However, highly reliable differences were observed regarding the parameters reflecting the speed of response: For the time of the mouse button press (see figure 2B), *t*(19)=−9.977, *p*<0.001, d_z_=−2.231, and to an even larger extent for movement onset (see figure 2C), *t*(19)=−12.48, *p*<0.001, d_z_=−2.791, the retro-cue condition revealed faster responses compared to the early probe condition.

**Figure 2.**
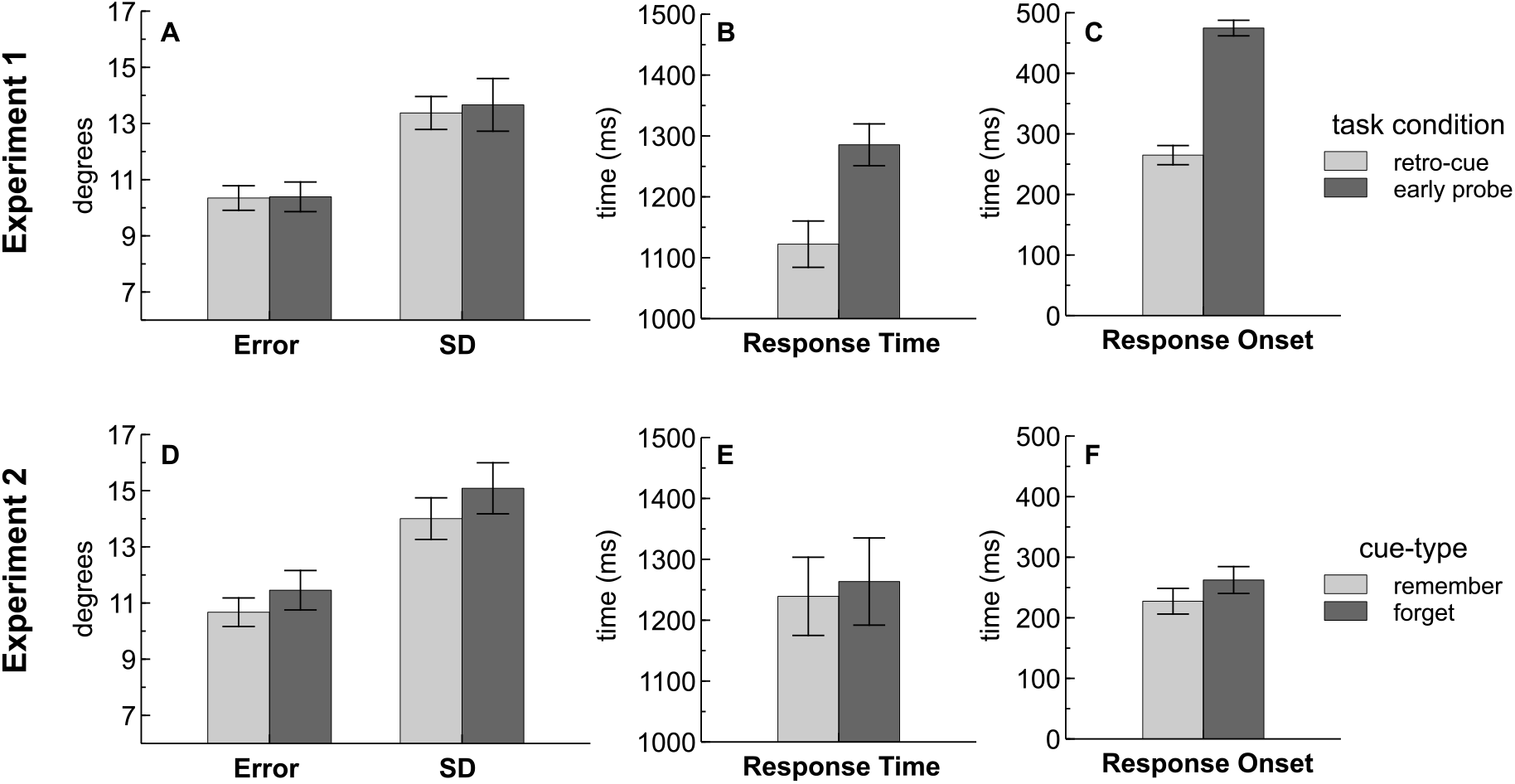
Behavioral data of Experiment 1 (**2A-C**) and Experiment 2 (**2D-F**). Task accuracy was defined as the raw angular error between the cued item from the memory array and the adjusted orientation. No difference in performance between the retro-cue and early probe condition was shown on the level of the raw error and the standard deviation (SD) of the raw error (**2A**). However, participants were faster in task completion (i.e. time until computer mouse button press; **2B**) and in starting the orientation adjustment (i.e. response onset time; **2C**) in the retro-cue compared to the early probe condition. In Experiment 2, task accuracy was higher for remember vs. forget retro-cues (**2D**). Also response onset time was faster following a remember compared to a forget cue (**2F**). Error bars reflect the standard error of the mean.

In addition, the continuous performance response allowed for fitting a mixture model on the data that divided the response distribution into four parameters (Bays, Catalao, & Husain, 2009): Parameter Kappa (k) reflects the precision of the recalled working memory representation independent from the item focused for recall. Parameters pT, pN and pU refer to the probabilities to report the target item (pT), the non-target item (pN) and the probability to report at random (pU). There was a trend toward a lower probability to report at random (pU) in the retro-cue compared to the early probe condition, *t*(19)=−1.726, *p*=0.101, d_z_=−0.386. Parameter pT, *t*(19)=1.571, *p*=0.133, d_z_=0.351, and also the remaining parameters did not reveal a difference between these conditions (all p-values > 0.56). Overall, these findings point toward behavioral benefits of the retro-cue that manifest in the speed of response but not in response accuracy.

#### 3.1.2. Experiment 2

The same behavioral parameters as in Experiment 1 were analyzed and within-subject *t*-tests were used to test for performance differences between the remember and forget cue conditions (two sided). Only trials with a mouse button press were included, with a rate of button presses across trials of 99.32 % on average (SD=0.97 %). Behavioral performance differed between the remember and forget cue conditions regarding the raw error, *t*(15)=−2.355, *p*<0.05, d_z_=−0.539, and SD of the raw error, *t*(15)=−2.1, *p*=0.053, d_z_=−0.525. Working memory accuracy was slightly increased following a remember compared to a forget cue (see figure 2D). Reliable differences were also observed regarding mouse movement onset (see figure 2F), *t*(15)=−2.655, *p*<0.05, d_z_=−0.664, with faster onsets in the remember cue blocks. There was no difference in response times (i.e., time point of button presses) between the remember and forget cues, *t*(15)=−0.574, *p*=0.575, d_z_=−0.143. Also the parameters of the mixture model did not reveal reliable differences between these conditions (all p-values > 0.13). Overall, results point toward higher performance in the working memory task when a remember compared to a forget retro-cue was presented.

### 3.2. EEG time-frequency analyses

Spectral power or event-related spectral perturbation (ERSP; Delorme & Makeig, 2004) was computed by convolving three-cycle complex Morlet wavelets with each epoch of the EEG data. Epochs consisted of 200 time points from −1000 to 3000 ms referred to the memory array and frequencies ranged from 4 to 30 Hz in 52 logarithmic steps. The number of cycles used for the wavelets increased half as fast as the number of cycles in the respective fast-fourier transformation (FFT). This resulted in 3-cycle wavelets used for 4 Hz and 11.25-cycle wavelets used for 30 Hz. Channels rejected during EEG data preprocessing were recomputed by means of spherical channel interpolation included in EEGLAB prior to these analyses.

#### 3.2.1. Experiment 1

The posterior lateral electrodes PO7/8, PO3/4, P7/8 and P5/6 were considered for further analyses. As it was not possible to define the analysis time windows and frequencies of interest a priori, we first averaged across the retro-cue and the early probe conditions and then calculated the contralateral vs. ipsilateral portions of the ERSPs. For trials with a lateralized target item, we averaged across left-sided channels and trials with cued memory items on the right side and right-sided channels and cued items on the left side (i.e., contralateral). The ipsilateral portion was accordingly calculated by averaging across trials with cued memory items on the right side and right-sided electrodes and cued memory items on the left side and left-sided electrodes. The same procedure was run based on the location of the non-cued item in the distractor lateralized condition. Attentional modulations in the ERSPs were then assessed by comparing the contralateral minus ipsilateral difference by means of within-subject *t*-tests between the target lateralized and distractor lateralized conditions for each ERSP data point. These tests were corrected for multiple comparisons by means of the false discovery rate (FDR) procedure (Benjamini & Hochberg, 1995). As shown in figure 3, these analyses revealed a significant cluster with increased contralateral vs. ipsilateral spectral power for the distractor lateralized compared to the target lateralized condition. The cluster appeared in the alpha frequency range (10-12 Hz) at about 430 ms to 600 ms following the retro-cue or probe display. Further analyses were based on these time and frequency ranges. An ANOVA was run including the within-subject factors *asymmetry* (contralateral vs. ipsilateral), *target position* (distractor lateral vs. target lateral) and *cue-type* (retro-cue vs. early probe). In both Experiment 1 and Experiment 2 (see below), post-hoc comparisons were based on the Tukey’s Honest Significant Difference procedure implemented in the Statistics and Machine Learning Toolbox of MATLAB®. There was a main effect of *cue-type*, F(1,19)=32.456, *p*<0.001, η_p_^2^=0.631, indicating a higher alpha power suppression over posterior sites for the early probe compared to the retro-cue condition. While the main effects of target position, F>1, and asymmetry, F(1,19)=2.013, *p*=0.172, η_p_^2^=0.096, were not significant, there was a reliable interaction of these factors, F(1,19)=37.397, *p*<0.001, η_p_^2^=0.663. This interaction was based on a contralateral vs. ipsilateral increase in alpha power for the distractor lateralized condition (M_diff_=0.291 dB, SE_diff_=0.078 dB, *p*=0.001) and the absence of a reliable asymmetry effect in the target lateralized condition, (M_diff_=−0.094 dB, SE_diff_=0.075 dB, *p*=0.225) (see figure 3). The retro-cue and the early probe conditions did not differ regarding this interaction of *asymmetry* and *target position*, F(1,19)=1.167, *p*=0.293, η_p_^2^=0.058. Overall, these results indicate that the lateralization of posterior alpha power as a marker of retroactive attentional orienting is closely related to the handling of the non-cued information.

**Figure 3.**
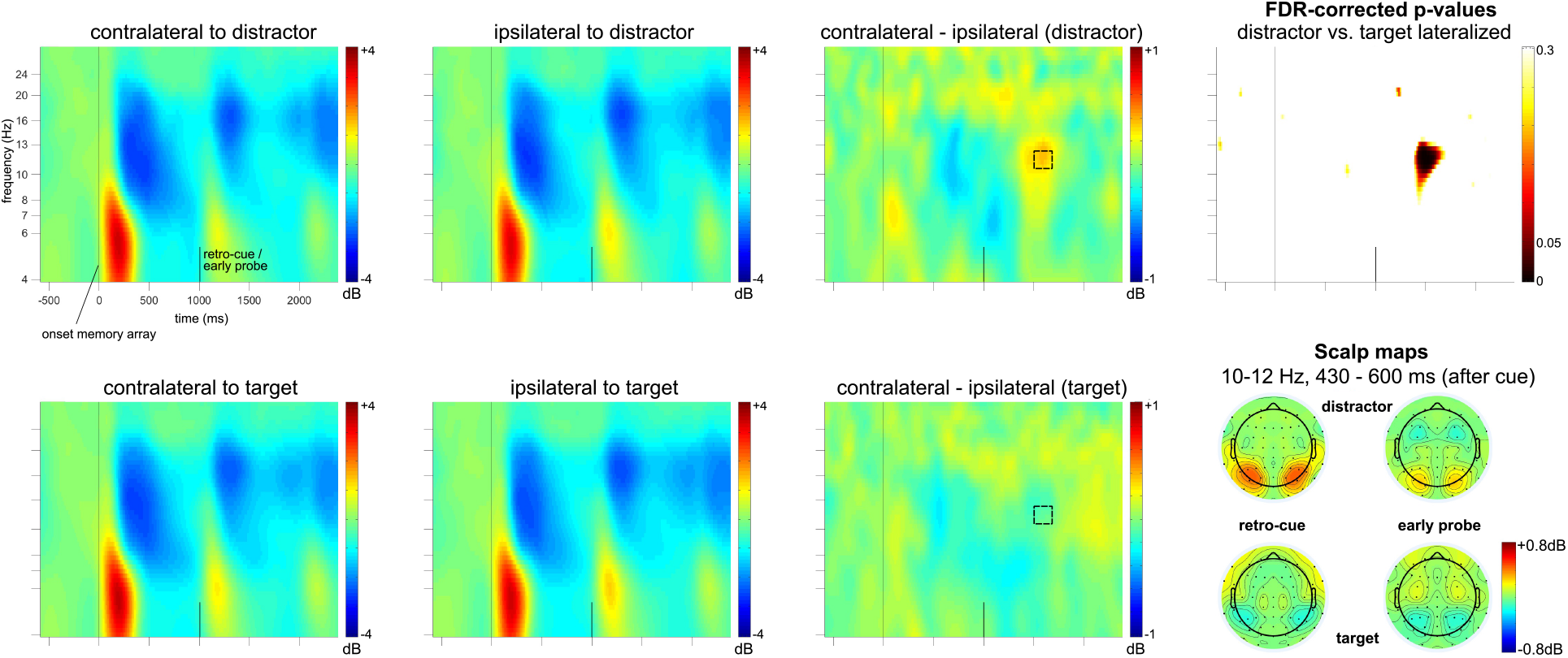
Lateralized effects in posterior ERSPs for Experiment 1. The oscillatory responses following the memory array (vertical line at zero) are shown contralateral vs. ipsilateral to a distractor (upper row) and target item (lower row) at a posterior lateral electrode cluster (PO7/8, PO3/4, P7/8, P5/6). ERSPs were averaged across the retro-cue and early probe conditions. The contralateral minus ipsilateral ERSP difference as well as the FDR corrected t-tests on the comparison of the target and distractor lateralized conditions indicated an effect in higher alpha frequency range at about 430 to 600 ms following the retro-cue and early probe displays. As also indicated in the respective contralateral minus ipsilateral difference topographies, this effect was related to higher alpha power contralateral compared to ipsilateral to the lateralized distractors.

#### 3.2.2. Experiment 2

We further intended to investigate if the posterior alpha asymmetries following retro-cues to lateralized targets or distractors differed between remember and forget cues. At first, analyses were kept comparable to Experiment 1. Analyses were based on the average signal of the posterior lateral electrodes PO7/8, PO3/4, P7/8 and P5/6. We averaged across the remember and forget cue conditions and then calculated the contralateral vs. ipsilateral portions of the ERSPs (see above). Attentional modulations in the ERSPs were measured by comparing the contralateral minus ipsilateral difference by means of within-subject *t*-tests (FDR corrected) between the target lateralized and distractor lateralized conditions for each ERSP data point. As shown in figure 4, these analyses revealed a significant cluster with increased contralateral vs. ipsilateral spectral power for the distractor lateralized compared to the target lateralized condition. The cluster appeared in the alpha frequency range (8-11 Hz) at about 600 ms to 730 ms following the retro-cue or probe display. Further analyses were based on these time and frequency ranges. We run an ANOVA with the within-subject factors *asymmetry* (contralateral vs. ipsilateral), *target position* (distractor lateral vs. target lateral) and *cue-type* (remember vs. forget cue).

This analysis only indicated a highly significant *target position* by *asymmetry* interaction, F(1,15)=26.118, *p*<0.001, η_p_^2^=0.635. While the distractor lateralized condition led to a contralateral vs. ipsilateral increase in posterior alpha power (M_diff_=0.293 dB, SE_diff_=0.08 dB, *p*=0.002), alpha power was decreased contralateral compared to ipsilateral to a lateralized target (M_diff_=−0.319 dB, SE_diff_=0.092 dB, *p*=0.004). All other interactions and the main effects of *cue-type, target position* and *asymmetry* failed to reach statistical significance (all F-values < 1). Retroactive attentional sub-processes did thus not differ between the remember and forget cues regarding the amplitude of posterior alpha power asymmetry (see also topographies in figure 4).

**Figure 4.**
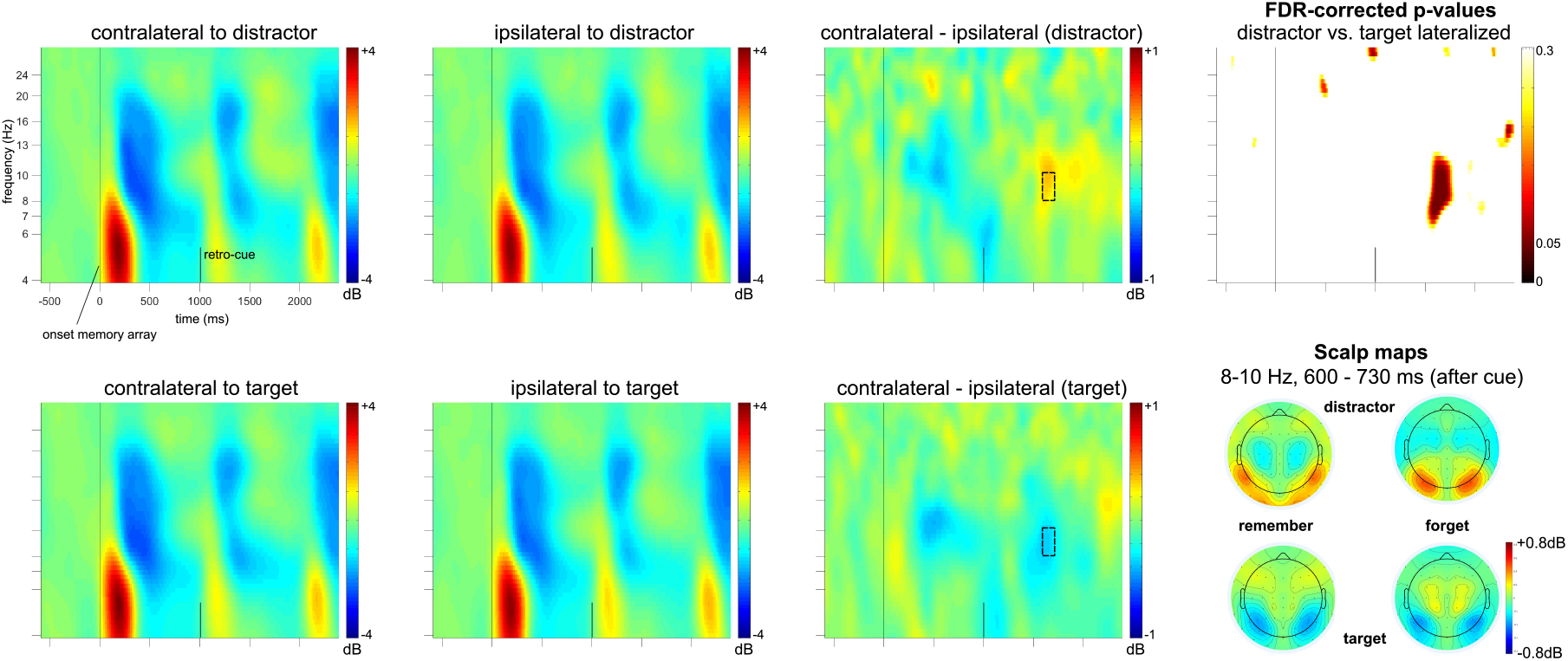
Lateralized effects in posterior ERSPs for Experiment 2. The oscillatory responses following the memory array (vertical line at zero), averaged across the remember and forget cue conditions, are shown contralateral vs. ipsilateral to a distractor (upper row) and target item (lower row) at a posterior lateral electrode cluster (PO7/8, PO3/4, P7/8, P5/6). The contralateral minus ipsilateral ERSP difference as well as the FDR corrected t-tests on the comparison of the target and distractor lateralized conditions indicated an effect in alpha frequency range at about 600 to 730 ms following the remember and forget retro-cues. As also indicated in the respective contralateral minus ipsilateral difference topographies for the remember vs. forget conditions, this effect was related both to target enhancement and distractor suppression.

However, differences in these cue-types might also appear in the latency of retroactive attentional orienting reflected in posterior alpha asymmetries. The latency of the alpha asymmetry was measured based on the contralateral and ipsilateral portions of the ERSPs based on electrodes PO7/8, PO3/4, P7/8 and P5/6 (see above). We chose an interval from 200 ms to 1000 ms following the retro-cue for further analyses. A start at 200 ms was chosen because attentional selection is unlikely to be reflected already in early time windows associated with sensory processing of the cue. The end at 1000 ms was chosen because it was also the onset of the probe display. Afterwards, we measured the negative (target lateral) and positive area (distractor lateral) under the contralateral minus ipsilateral difference wave within each experimental condition and for each participant. The latency of the alpha asymmetry was defined as the point in time when the area under the curve reached 50% of its total value. An ANOVA with the within-subject factors *target position* (distractor lateral vs. target lateral) and *cue-type* (remember vs. forget cue) was run based on the latency values. We observed a reliable interaction between these two factors, F(1,15)=10.45, *p*<0.01, η_p_^2^=0.411. The main effects of *target position* and *cue-type* were not significant (*all F*-values < 1). While there was an earlier alpha asymmetry latency for forget vs. remember cues when the distractor was lateralized (M_diff_=−118.31 ms, SE_diff_=65.482 ms, *p*=0.091), the asymmetry was delayed following forget vs. remember cues when the target was presented lateralized (M_diff_=95 ms, SE_diff_=41.281 ms, *p*=0.036) (see figure 5). This indicates that the latency of retroactive attentional sub-processes depended on the cue-type, with remember cues facilitating target enhancement and forget cue facilitating distractor inhibition processes. As a side note, a comparable interaction effect was also observed when considering the time point of 20% total area under the curve as a marker for the onset of the posterior alpha asymmetry, F(1,15)=9.61, *p*<0.01, η_p_^2^=0.39, but not when the time point of 80% area under the curve was used, F(1,15)=0.222, *p*=0.644, η_p_^2^=0.015. This indicates that the respective modulation was related to the early phases of retroactive attentional selection.

**Figure 5.**
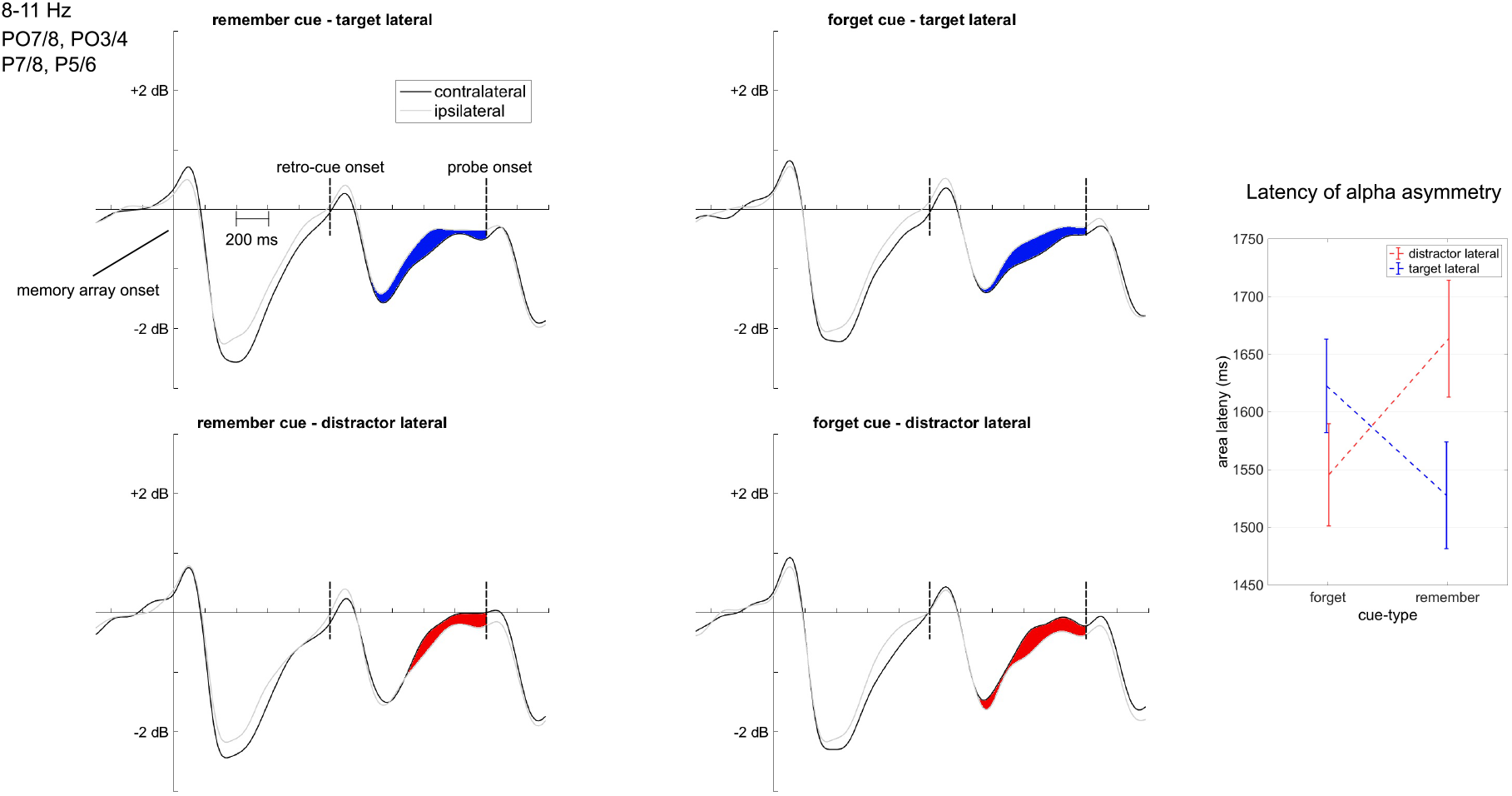
Posterior contralateral vs. ipsilateral alpha power (8-11 Hz) for Experiment 2. The black line depicts the contralateral ERSPs at the posterior lateral electrode cluster (PO7/8, PO3/4, P7/8, P5/6), while the gray line represented the ipsilateral portion of the signal. Contralateral vs. ipsilateral lineplots are shown for the remember vs. forget and the distractor vs. target lateralized conditions. The blue areas stand for a stronger contralateral vs. ipsilateral alpha suppression (i.e., ‘negative area‘). The red areas depict increased alpha power contralateral to the distractor (i.e., ‘positive area’). The interaction plot on the right provides the time points when 50% of the positive or negative areas under the contra-minus ipsilateral difference curves were reached in the remember vs. forget conditions and target vs. distractor lateralized conditions. Error bars reflect the standard error of the mean.

## 4. Discussion

The current study investigated the contribution of target enhancement and distractor suppression mechanisms to retroactive attentional orienting. In two delayed estimation working memory paradigms, a retro-cue or memory probe indicated either a lateralized item (i.e., target lateralized condition) or non-lateralized item from a bilateral memory array (i.e., distractor lateralized condition) for report. Results suggest that both attentional sub-processes play an important role in retroactive attentional orienting and allow for the interpretation of both mechanisms as active cognitive processes.

On behavioral level, retro-cues led to a performance benefit compared to the early probe condition (see figure 2). While there was no reliable difference between these conditions based on the accuracy and mixture model parameters, responses were faster in the retro-cue compared to the early probe condition. This effect was observed both for the response onset time (i.e. the time between probe onset and the first computer mouse movement) and the time required to complete the rotation movement (i.e. the button press). This suggests that retro-cues led to the faster completion of response preparation, a finding that is in line with earlier observations on a head-start of item retrieval and response planning mechanisms following a reliable retro-cue (Myers, Stokes, & Nobre, 2017; Schneider, Barth, & Wascher, 2017; Souza, Rerko, & Oberauer, 2016). While in the early probe condition the retrieval of the cued item and the processing of the probe had to proceed simultaneously, the retro-cue allowed to select the cued mental representation ahead of the probe information and already transfer it into a response-oriented representational state (i.e. response planning). Prior studies also found a retro-cue benefit on accuracy level compared to an early probe condition (e.g., Murray, Nobre, Clark, Cravo, & Stokes, 2013; Souza, Rerko, Lin, & Oberauer, 2014). However, in these studies, the set size of the initial memory array containing differently colored items was up to eight stimuli. Souza and colleagues (2014) showed that increasing the set size of the memory array from one (no retro-cue effect expected) to four items led to an increase of the retro-cue benefit on the level of accuracy compared to an early probe condition and remained stable from set sizes four to eight. It can thus be assumed that the focusing on one out of two memory representations in the current study did not suffice for revealing reliable retro-cue benefits on the level of accuracy.

The separation of attentional sub-processes during retroactive attentional orienting was based on lateralized effects in the posterior alpha response following the retro-cues and probes. We thereby intended to answer the question whether attentional selection within working memory contents included the inhibition of the non-cued representations, or whether it was exclusively related to the transfer of the cued representation into a higher-level representational state. Several studies investigated the lateralization of posterior alpha power as an indicator for retroactive attentional orienting (e.g., Myers et al., 2015; Poch, Campo, & Barnes, 2014; Poch, Capilla, et al., 2017; Poch, Carretie, et al., 2017; Schneider et al., 2015, 2016). Comparable to the current experiment, Poch, Capilla and colleagues (2017) presented endogenous color retro-cues toward memory items presented on the left or right side of fixation and revealed a higher decrease in alpha power contralateral compared to ipsilateral to the cued contents. This effect was not sustained throughout the delay period and was thus associated with a temporary attentional process rather than working memory storage. The lateralization of alpha power in the context of attentional orienting during perception was interpreted as the consequence of an inhibitory control process setting the neural population processing irrelevant signals into a kind of ‘idling state’. This state goes along with an increase in alpha power contralateral to irrelevant or distracting information (Handel, Haarmeier, & Jensen, 2011; Kelly, Lalor, Reilly, & Foxe, 2006; Klimesch, Sauseng, & Hanslmayr, 2007; Rihs, Michel, & Thut, 2009; Sauseng et al., 2005). While earlier studies also discussed the possibility that inhibitory mechanisms contribute to retroactive attentional orienting (Poch, Carretie, et al., 2017), the majority of investigations in the field of working memory rather favored the notion that alpha suppression is related to target processing (Fukuda, Mance, & Vogel, 2015; Myers et al., 2015; Schneider et al., 2016).

In the current study, we approached these inconsistencies by strictly separating target enhancement from distractor inhibition processes based on the spatial layout of the memory array. In Experiment 1, there was an increase in alpha power contralateral compared to ipsilateral to a non-cued item, but no alpha asymmetry when the target item was presented lateralized. There are two possible ways this dealing with the irrelevant item or distractor might proceed: A lateralization of posterior alpha power might reflect a control process for actively inhibiting the irrelevant mental representation or it might be the consequence of an automatic inhibition of the distractor item by means of the attentional template for the target (Desimone, 1998; Desimone & Duncan, 1995). In both cases, retroactive attentional orienting should lead to a change of the irrelevant representation into a more passive and fragile short-term memory state (e.g., Sligte, Scholte, & Lamme, 2008; Vandenbroucke, Sligte, de Vries, Cohen, & Lamme, 2015; Vandenbroucke, Sligte, & Lamme, 2011), while leaving the target item within the focus of attention in working memory. However, when comparing Experiment 1 and 2, it becomes obvious that it is problematic to interpret the lack of alpha asymmetry following a retro-cue toward a lateralized target as a sign for the dominance of distractor inhibition in retroactive attentional orienting. Experiment 2 revealed a clear contralateral suppression of posterior alpha power following a cue toward a lateralized target (see figure 4). As also Experiment 1 indicated a minor asymmetry in the target lateral condition on descriptive level, we would thus conclude that both target enhancement and distractor inhibition contribute to retroactive attentional orienting.

By also presenting cues indicating which representation to remember or forget, we further investigated if target enhancement and distractor suppression can be interpreted as separate active processes (cf., Williams, Hong, Kang, Carlisle, & Woodman, 2013; Williams & Woodman, 2012). While we usually think about the orienting of attention toward relevant information as a deliberate and active process, there is a logical problem regarding active inhibition: How can we ‘think about’ or process something and thereby inhibit it? Or in terms of a memory representation, how can the processing of any content lead to forgetting? There is, however, evidence that cuing an irrelevant item prior to visual search leads to faster responses to the target. While this effect might be interpreted by a reduction of ambiguity about the target, cuing distractor locations also reduced the distractors’ automatic interference with target processing in a flanker-like experiment (Munneke, Van der Stigchel, & Theeuwes, 2008), supporting the active inhibition hypothesis. Referred to the current paradigm, cuing a no longer relevant item might result in the update of the attentional template to only include the target feature. In this view, neural correlates of distractor inhibition would be related to an automatic lateral inhibition caused by the refined attentional template (Desimone, 1998; Desimone & Duncan, 1995). A remember cue would thus be used to directly update the attentional template, but a forget cue would involve only indirect information about the target. Excitatory and inhibitory attentional sub-processes should thus be facilitated following a remember compared to a forget cue, or should at least not differ between these conditions provided that the reframing of the forget cue proceeded at low costs. Alternatively, selective forgetting in terms of the removal of irrelevant representations from working memory might be involved, as for example proposed by the SOB-CS model of working memory processes in the complex span task (Oberauer, Lewandowsky, Farrell, Jarrold, & Greaves, 2012). Such a mechanism might be directly triggered by a cue to forget a certain working memory representation. On behavioral level, both the remember and forget cues enabled adequate task performance. However, the remember cue was associated with higher performance based on both accuracy and response time parameters compared to the forget cue, suggesting that retrieval of the relevant memory item was more efficient with remember compared to forget cues. However, intriguingly, the electrophysiological results did not reveal a comparable main effect in favor of the remember cues (see figure 5): Distractor inhibition proceeded faster when the distractor was cued, but target selection was faster following a remember compared to a forget cue. It is not possible to interpret these findings in the light of target selection and automatic lateral inhibition. We would thus suggest that both distractor inhibition and target selection within visuo-spatial working memory are separable active processes.

In summary, the current study provided important insights into the attentional sub-processes underlying the updating of visuo-spatial working memory representations. Retro-cues allowed for target selection and response planning ahead of the presentation of the memory probe, thereby leading to faster responses in a delayed estimation working memory task compared to the early probe condition. Attentional selection based on the retro-cues or probes was investigated by means of lateralized effects in oscillatory power. We observed an increase in posterior alpha power contralateral compared to ipsilateral to the distractor item reflecting inhibition of the no longer relevant representation. Additionally, there was contralateral suppression of posterior alpha power when the target was presented lateralized. It can thus be assumed that the alpha lateralization following retro-cues in earlier studies (e.g., Myers et al., 2015; Poch et al., 2014; Poch, Capilla, et al., 2017; Poch, Carretie, et al., 2017; Schneider et al., 2015, 2016) was related to both an enhancement of the relevant mental representations and to changes in the representational state of the non-cued working memory contents. Interestingly, cues indicating the item to remember and to forget led to distinct differences in these attentional processes. While retroactive attentional selection of the target was faster following a remember compared to a forget cue, the opposite pattern appeared for distractor inhibition. This proposes that retroactive attentional orienting proceeds based on two separate cognitive sub-processes related to the target and to the no longer relevant mental representations.

## 5. Acknowledgements

The study was supported by a grant from the German Research Foundation (DFG Grant No. SCHN 1450/1-1) to Daniel Schneider.

